# Generalizing the response and satisfaction threshold models for multiple task types: A maximal entropy approach

**DOI:** 10.1101/2024.04.26.591219

**Authors:** C.M. Lynch, T.P. Pavlic

## Abstract

Response threshold models are often used to test hypotheses about division of labor in social-insect colonies. Each worker’s probability to engage in a task rapidly increases when a cue associated with task demand crosses some “response threshold.” Threshold variability across workers generates an emergent division of labor that is consistent over time and flexibly adaptive to increasing demands, which allows for testable predictions about the shape of hypothetical response-threshold distributions. Although there are myriad different task types in a social-insect colony, the classical response-threshold model is built to understand variability in response to a single type of task. As such, it does not immediately allow for testing predictions about how different workers prioritize different task types or how demand for some tasks interferes with responding to demand for others. To rectify this, we propose a multi-task generalization that degenerates into the standard model for a single task. We replace the classical Hill response probability with a model that draws worker choices from a Boltzmann distribution, which is an approach inspired by multi-class machine learning.

## 1. Introduction

The response threshold concept is an extremely parsimonious explanation of social insect behavior, combining explanations of plasticity and specialization into a single mechanistic framework (Beshers & Fewell, 2001). Under this paradigm, individual workers in a colony intrinsically differ in the cue level at which they start working, and by performing the task associated with that cue, they reduce the cue and therefore also reduce the probability that this task will be performed by others (Jeanson & Weidenmüller, 2014). Models have shown that response thresholds can increase the flexibility and robustness of a colony in response to environmental perturbations (Calabi, 1988; Camazine et al., 2001) and it can act as a proximate mechanism behind division of labor (Beshers et al., 1999). Empirically, response thresholds have also captured social behaviors related to division of labor (Fewell & Page, 1999; Pankiw & Page, 2000; Holbrook et al., 2011; Holbrook et al., 2013; Perez et al., 2013; Brahma et al., 2018; Rajagopal et al., 2022). Despite these successes, response threshold models remain limited in their ability to capture many ecologically relevant behaviors at the heart of division of labor and other phenomena (Garrison et al., 2018; Leitner et al., 2019; Ulrich et al., 2021).

Response thresholds are traditionally modeled with a Hill function, which takes a task-associated cue as an input and outputs an s-shaped curve which gives the probability of starting a single task (Bonabeau et al., 1996). Models using this function successfully predicted an uptick in division of labor in larger harvester ant colonies (Jeanson et al., 2007; Holbrook et al., 2011) as well as increased homeostasis for small clonal raider ant groups (Ulrich et al., 2018). It has also been used to solve task allocation problems in swarm robotics (Wu et al., 2018; Jiang et al., 2020). However, this function is limited in two respects.

1) The Hill function breaks down for negative task-associated cues. Traditionally, task-associated cues are thought to be physical cues that can only take on positive real values (Bonabeau et al., 1996), or the probability of starting a task must be set to 0 with a piecewise function if the cue becomes negative (Yang et al., 2009). Modeling them this way works for cases where the social insect colony wants to either minimize the cue (such as with brood pheromone; Pankiw et al., 1998) or maximize it (Gove et al., 2009). However, there are cases where an intermediate value of the cue is preferred, such as with nest temperature (Weidenmüller, 2004; Gove et al., 2009). In these cases, it may make sense to model the task-associated cue not as the raw cue, but rather as a distance from an objective, which could take on a negative value. Having a low temperature - or a negative objective value - may make the probability of fanning increasingly small while the probability of incubating brood increasingly large (O’Donnell & Foster, 2001).

2) The Hill function has difficulty incorporating multiple tasks. Naively applying this function to multiple tasks simultaneously without a normalization protocol can result in probabilities that sum to a value greater than one. Multiple approaches have been proposed to solve this issue while still using the Hill function. Some assume that there is only one task (Bonabeau et al., 1996; Arcuri & Lanchier, 2017; Feng et al., 2021), others allow ants to randomly encounter tasks one at a time (Jeanson et al., 2007; Ulrich et al., 2018; Lin, 2021; Ulrich et al., 2021) or perform the task of greatest need which exceeds the individual’s threshold (Gove et al., 2009), and finally some calculate relative cues and thresholds so that all tasks can be experienced concurrently (Wu et al., 2018; Jiang et al., 2020; Lynch et al., *in submission*). The assumption that there is only one task is likely problematic in many cases, as real social insects perform multiple tasks, and variable task numbers can influence model performance (Dornhaus et al., 2019). Random encounter models can be biologically reasonable within specific contexts, such as within nests where cues may be locally isolated (Ravary et al., 2007; Jeanson & Lachaud, 2015). However, this assumption is not always met, as social insects can have more open nests (Tschinkel, 2005) which can allow them to encounter multiple pheromones at the same time. They can then differentiate these signals with different types of sensilla (Dussutour et al., 2009). Reprocessing a task-associated cue as a relative value to other cues can avoid the assumptions of the previous two approaches, however as we will show, this approach tends to uniformly distribute the probabilities of performing different tasks, which may or may not be a desirable property.

A special form of a Boltzmann sampler can be used to resolve both these issues. Boltzmann samplers are designed to sample from different types of combinatorial structures, which will have a certain size depending on how many different combinations of elements there are within that structure (Duchon et al., 2004). In the field of statistical mechanics, physicists are interested in describing the distribution of particles in different states. Higher-energy states are less likely to occur than lower-energy states, as there are fewer possible combinations of microscopic configurations that result from that energy. When one equates energy with the size of combinatorial structure while also including the temperature of the system, the Boltzmann sampler returns the Boltzmann distribution, the maximal entropy probability distribution across these energy states. As this distribution includes a partition function which guarantees that the sum of the probabilities will be 1, this distribution has applications in many other fields. Most famously, it is known as the softmax function in data science, where it is used as the last activation function of a neural network. For neural networks, the softmax function normalizes the output of the algorithm so that the scores associated with various classes can be interpreted as probabilities. This function has also been used to find the probability of performing different actions in the field of reinforcement learning (Sutton & Barto, 1998), and so it seems like a natural candidate for modeling response thresholds.

In the case of social insects, the difference between the task-associated and the response threshold cue is analogous to the energy of the system (or the scores of a neural network). If we further equate the inverse of the temperature with the steepness of the probability curve (the Hill coefficient), then the Boltzmann sampler gives a mathematically tractable way of normalizing the probabilities of performing different tasks that can allow for both positive and negative task-associated cues. This also offers an interpretation of how a colony integrates multiple task-associated cues in a way similar to that of physical particles, but it allows these particles some degree of volition by giving each their own threshold. The Boltzmann sampler also has a number of other desirable properties that make it a natural extension of the Hill function. The function has point symmetry in the single task case, the steepness of the function only depends on a single parameter, the function is scale-free, and it emphasizes the effect of the thresholds on an individual social insect’s behavior. It can also be easily modified to incorporate other phenomena as well, including task completion cues and satisfaction thresholds (Lynch et al., *in submission*). Together, these properties ease the process of model construction, allowing for scenarios that would not have been possible to model using traditional Hill functions.

## 2. Building Threshold Models with a Boltzmann Sampler

### 2.1. Single task response thresholds models can be derived from a softmax function

Response thresholds are typically modeled probabilistically. That is, different values of the task-associated cue (*x*) yield different probabilities of a response *P*(*x*, *θ*; *k*). This is usually written as a Hill function:

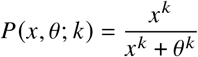

which generates an s-shaped probability curve. Here, *θ* is the inflection point (the response threshold) of the curve while *k* (the Hill coefficient) determines the steepness of the curve. As *k* approaches infinity, the curve approaches a stepwise function, and can be interpreted as a measure of how deterministic the social insect’s choice is. However, as Kanakia et al. (2016) showed, the response threshold can be re-imagined as the logistic function *S*(*x*, *θ*; *k*):

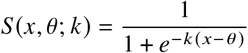

Here, the numerator sets the maximum level of the logistic, which we set to 1 as we interpret this output as a probability. *S*(*x*, *θ*; *k*), however, can only model 2 states: the worker is either performing the task or not performing it. If there are multiple potential outcomes, then a separate logistic function cannot be used for each task. As with the Hill function, the sums of the resulting probabilities could be greater than 1. We can instead derive the logistic function from a more general and slightly modified version of the softmax function:

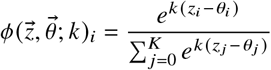

where 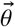 is the vector which contains the response thresholds for each task and 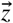 contains the task associated cues for *K* + 1 states for *K* tasks and an additional resting state. In this context, *k* is the inverse of temperature (and thus *k* ≥ 0) for both the Boltzmann distribution and the softmax function used in neural networks, which are themselves special cases of the Boltzmann sampler. When 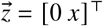 and 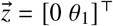 where 0 indicates that there is no response threshold for resting, and we solve for state 1, the softmax function simplifies to:

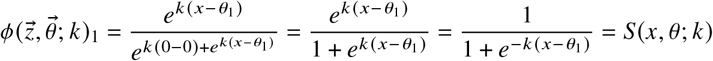

The sum of 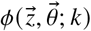 across *i* is guaranteed to be 1 as we are normalizing by the sum, so this function can be used to model a set of *K* response thresholds for an individual social insect.

### 2.2. Incorporating satisfaction thresholds and task completion cue

We can further generalize the softmax function by allowing it to incorporate satisfaction thresholds as well as task completion cues (Lynch et al., *in submission*). To review, response threshold models usually assume that the signals that determine a need for a task increase over time, and can therefore be interpreted as an ‘on’ signal called a task demand cue. Satisfaction thresholds determine the probability of stopping a task rather than starting a task. Therefore, the probability of stopping tasks should decrease as the task demand cue increases. However, if the signal decreases over time rather than increases, then it can be interpreted as an off signal and is called a task completion cue. The probability of starting a task should decrease with higher task completion cues, and satisfaction thresholds should increase. We therefore need a function that can create s-shaped curves with a negative as well as a positive first derivative.

To do this, we can introduce two binary variables *α, β* ∈ {−1, 1}. *α* is encoded such that 1 represents a task demand cue and −1 represents task completion. *β* is similar where 1 represents a response threshold model and −1 represents a satisfaction threshold model (Fig. 1). Multiplying these variables determines the sign of the exponent, which in turn determines the slope of the resulting probability curves. For response thresholds, these curves represent the probability of starting a task whereas for satisfaction thresholds these represent the probability of stopping a task. The general function can be written as:

**Fig. 1.**
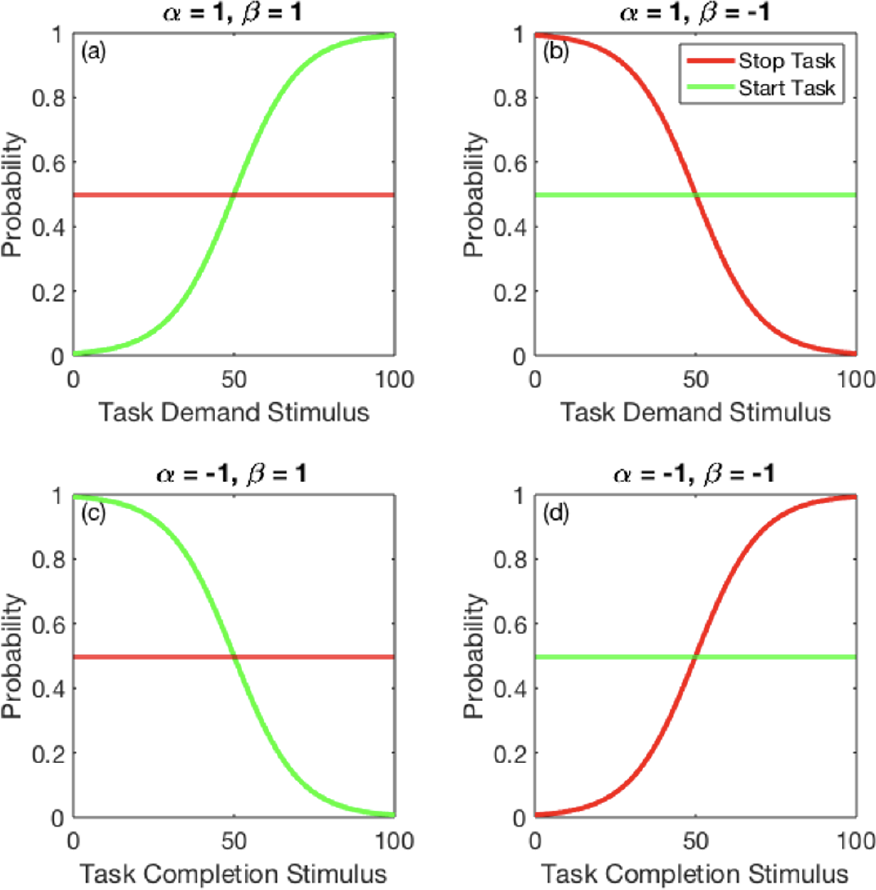
*ϕ* and *ϕ*′ functions when *θ* = 50, *k* = 0.1, *c* = 0.5, and *K* = 1. In each case, the logistic is generated from *ϕ* while the horizontal line is generated from *ϕ*′. Red lines show the probability of stopping a task, and the green lines show starting probabilities. Each panel shows the effect of different combinations of *α* and *β* on the logistic curve. (a) represents a response threshold model with a task demand cue, (b) satisfaction threshold model with task demand cue, (c) response threshold model with task completion cue, (d) satisfaction threshold with task completion cue.

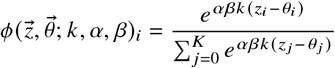

Note that if *α* × *β* = −1, and *K* = 1, then the first element of this function reduces to:

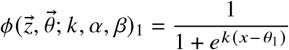

This function, like the probability function used in Lynch et al. (*in submission*) sums to unity for all x when added to the *α* × *β* = 1, version of the function, indicating that the two functions are complements of one another:

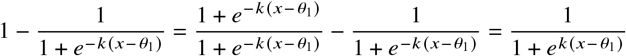

Response and satisfaction thresholds can also be incorporated into the same model of an ant to produce a composite threshold (Lynch et al., *in submission*). Response threshold models usually assume that the probability of stopping and starting (respectively) are constant (Bonabeau et al., 1996; Jeanson et al., 2007; Jiang et al., 2020). That is, the mirror of the *ϕ* function, *ϕ*′, is a constant 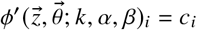 where 0 ≤ *c*_*i*_ ≤ 1. It is also possible to force both the starting and stopping probabilities to be dependent on the task-associated cue, which would be a composite threshold model. In this case, the mirror function would be:

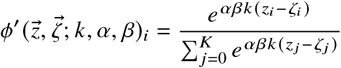

where the vector 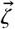 contains the thresholds for the opposite operation, depending on how one defines the primary function. For example, if *ϕ* determines the probability of starting a task, 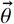 would contain response thresholds while *ϕ*′ would determine the probability of stopping the task and 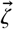 would contain satisfaction thresholds (Fig. 1).

## 3. Useful Properties of the Boltzmann Sampler

### 3.1. Softmax probabilities automatically sum to one

The softmax function allows for flexible model building. For instance, transitions between states can be effectively disallowed by making the threshold for a transition extremely high when *α* × *β* = 1, or extremely negative when *α* × *β* = −1. These transitions can also be used for continuous-time models or discrete-time models. Additionally, the partition function in the denominator will allow any distribution of thresholds can be used and the function will still sum to 1, regardless of how thresholds are distributed:

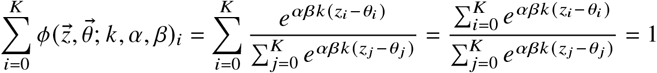

This result still holds even if *α, β*, and *k* are vectors. This means that using the softmax function makes it possible to model different types of tasks simultaneously. For instance, some tasks which are difficult for social insects to characterize could have a low value of *k*_*i*_, some may be modeled by a task completion cue, and yet others can have a response or satisfaction threshold.

The partition function also allows for any distribution of thresholds. For instance, a modeler can explore how threshold distributions may affect colony performance (Beshers & Fewell, 2001), as any real-valued threshold landscape will be possible using this sampler. Thresholds can change over time with reinforcement (response thresholds; Theraulaz et al., 1998; Giurfa & Sandoz, 2012) or desensitization (satisfaction thresholds; Maccaro et al., 2020; Lynch et al., *in submission*). They can even be correlated with one another via copulas regardless of one’s choice of threshold distribution (Bouyé et al., 2000). However, there is the issue of overfitting. Including too many higher-order correlational structures may allow a model to fit any dataset. The most parsimonious assumption then - that thresholds are independent of one another - may be the most advantageous in many situations. This partition function can also allow the steepness parameter to be a vector if tuning the stochasticity of different tasks is biologically meaningful or has some adaptive benefit. Finally, thresholds also change according to the behavioral context (Sazaki et al., 2014; Sadeh & Clopath, 2022; Moreno & Arenas, 2023).

### 3.2. Task associated cues can be negative for the softmax function

*P*(*x*, *θ*; *k*) breaks down when *x* < 0, as it is either undefined on the real number line for fractional values of *k* or tends to increase for whole numbers of *k*. This latter property can be problematic in cases where there is a negative task-associated cue. Conversely, the softmax function can easily incorporate negative distances without any of the issues that arise with the Hill function. To see this, we can evaluate the limit of the softmax function as *z*_*i*_ → −∞ for different signs of the exponent given different signs of the exponent:

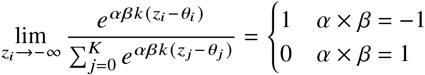

We can do the same for *z*_*i*_ → ∞:

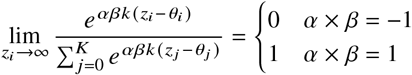

Thus, we can see that the function will always be constrained between 0 and 1 regardless of the sign of the exponent and whether *z*_*i*_ is positive or negative.

### 3.3. Probabilities curves around softmax threshold are symmetric

The function *S*(*x*, *θ*; *k*) has point symmetry around (*θ*, 1/2) such that *S*(*x*, *θ*; *k*) for *x* > *θ* is a 180^°^ rotation of *S*(*x*, *θ*; *k*) for *x* ≤ *θ*. This happens because *S*(*x*, *θ*; *k*) satisfies the condition 1 − *S*(*x*, *θ*; *k*) = *S*(−*x*, *θ*; *k*) where *θ* translates the inflection point (the point of symmetry) along the x-axis. The function *P*(*x*, *θ*; *k*) does not have this property. While *P*(0, *θ*; *k*) = 0, the function asymptotes in the positive x direction as:

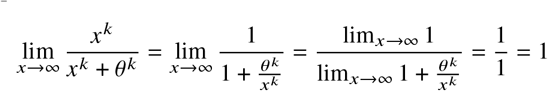

If *P*(*x*, *θ*; *k*) represents a response threshold, this means that a virtual ant is guaranteed to not perform a task when the task demand cue is 0, but is not guaranteed to start the task even if the cue is extremely high. Therefore, this function does not have point symmetry.

### 3.4. Steepness property dependent on only a single parameter for the softmax function

The steepness of the function Hill function *P*(*x*, *θ*; *k*) is purported to be controlled *k*, however this is decidedly not the case as the steepness of the function also depends on *θ* (Fig. 2). This is problematic for biological models, as this makes it more difficult to interpret the effects of each parameter in isolation of one another. For swarm roboticists, this is problematic as it makes it more difficult to control the exact shapes of these curves as neither parameter is truly free of the other. On the other hand, these parameters are largely independent for *S*(*x*, *θ*; *k*). We can show this by first taking the derivative of this function:

**Fig. 2.**
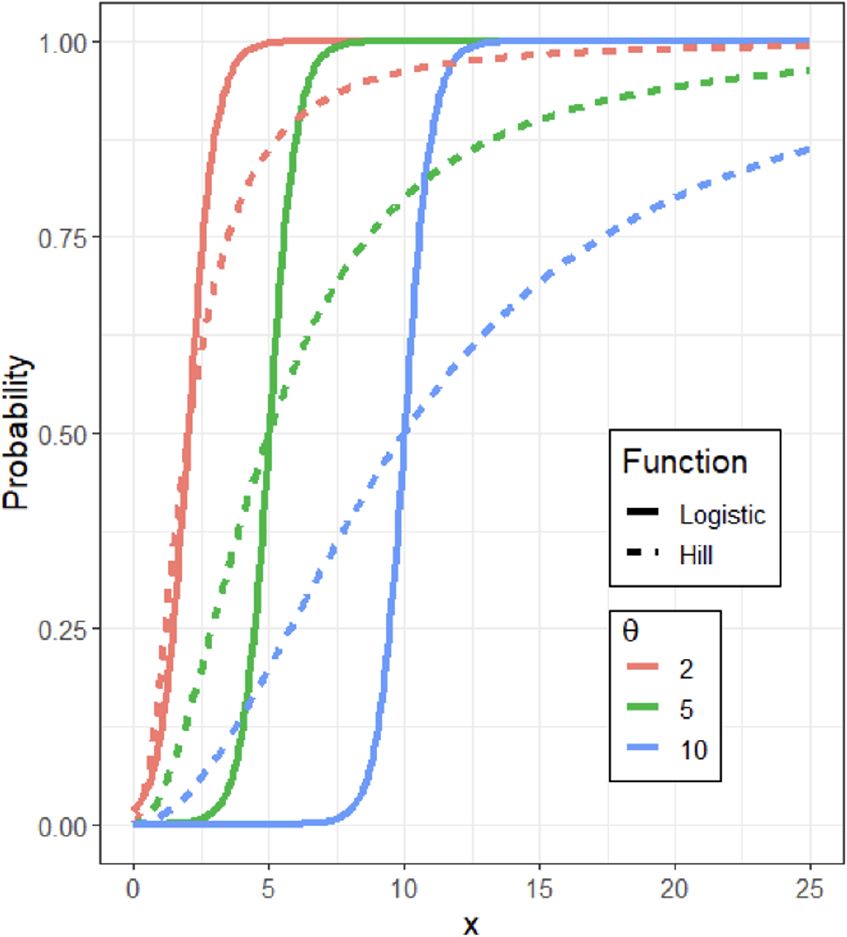
Response probability curves for Hill (dashed line) vs softmax functions (solid line) across different values of *θ* (color). In these plots, the steepness parameter *k* for both functions is set to 2. Note that the probability curves for the logistic tend to be parallel to one another despite variation in *θ*. The same is not true for the Hill function.

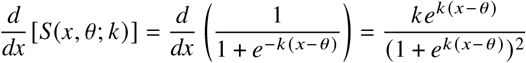

At the inflection point, *x* and *θ* cancel out each other and this expression simplifies to:

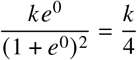

Thus, the steepness of *S*(*x*, *θ*; *k*) only depends on *k*. This is not the case for *P*(*x*, *θ*; *k*). Taking the derivative of this function yields:

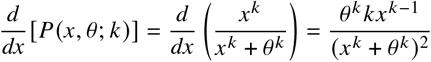

We can now evaluate the slope of *P*(*x*, *θ*; *k*) at the inflection point where *x* = *θ* = *ν*:

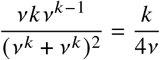

So we can see that as *θ* increases, the slope of *P*(*x*, *θ*; *k*) at the inflection point decreases, so slope does not purely depend on *k*.

This lack of dependence on the threshold also holds for the general, multi-task case for softmax. Let 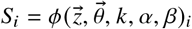 and *a* = *αβk*, then the partial derivative for the softmax function is:

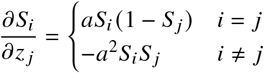

See supplemental information for derivation. In the *i* = *j* case:

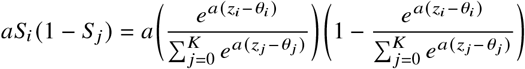

At the inflection point of all tasks, *z*_*i*_ = *θ*_*i*_ ∀ *i* = 1, 2, …, *K*, so:

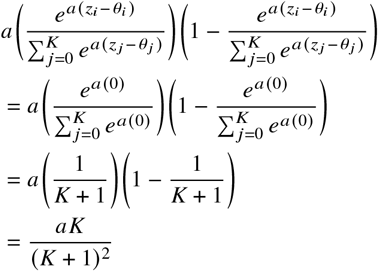

Performing this same procedure for *i* ≠ *j* yields −*a*^2^/(*K* + 1)^2^. As *α* and *β* can only equal −1 or 1, they can only determine the sign of the slope. Therefore, the steepness of the probability gradient of the threshold functions does not depend on *θ*, only on *k* and the number of states.

### 3.5. Scale-free nature of softmax function

The debate on how to integrate multiple cues is similar to the debate on how to convert the output of a neural network into a probability. The naive approach would be to use normalization, which would conserve the relative proportions of each score. Conversely, the softmax function down-weighs non-maximal values without eliminating them, increasing the probability that the highest score will be chosen in a process perhaps not dissimilar to that of a real neural system. One method of setting the probabilities to do tasks with response thresholds is similar in spirit to normalization. In Lynch et al. (*in submission*), the probability of performing a task for a response threshold task demand model is the response threshold multiplied by the bias *b*_*i*_ for that particular state *i*:

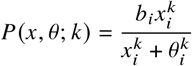

where *b*_*i*_ is the relative strength of a focal task-associated cue compared to all other cues:

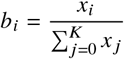

This term guarantees that the sum of probabilities across all tasks will be less than 1. Similar to normalization by sum, it will assign a probability of 0 to a task which has a task-associated cue of 0 (which may or may not be biologically feasible), it has trouble handling negative values, and when the thresholds for all tasks are equal, then the relative proportions of cues are correlated in the relative probability values. None of these properties are shared by the softmax function.

Consider the following numeric example. Here, we find the output of both the Hill and softmax functions given the vector [2 4 3]^⊤^. To isolate the effect of the function type, we set all *θ*_*i*_ = 1, and we set *k* to its typical value of 2. The first function is the Hill, and the latter the softmax (note that the output vector for the Hill does not sum to 1, as the remaining probability is allocated to the rest state. None of the task-associated cues were made to represent the rest state for the softmax function, so this output sums to 1. This difference is irrelevant for this demonstration):

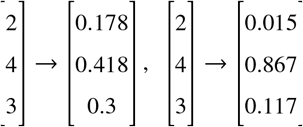

For the Hill function, we can see that the output probabilities are close to the bias for its corresponding cue (for example, 2/(2 + 4 + 3) ≈ 0.222 ≈ 0.178). In fact, as *k* increases, these probabilities will converge on the biases when *x*_*i*_ > *θ*_*i*_ or will go to 0 when x_*i*_ < *θ*_*i*_ or *b*_*i*_/2 when x_*i*_ = *θ*_*i*_. Conversely, the softmax function will disproportionately promote the highest cue, although lowering the response thresholds (or increasing satisfaction thresholds) of the other cues will mute this effect. Decreasing *k* can also soften the difference between these probabilities. Notably, the softmax function will retain these probabilities even if a constant is added to all the cues. Here, we add 100 to the input vector:

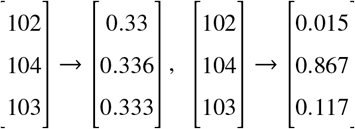

This occurs because when you add any constant ∀ *r* > 0:

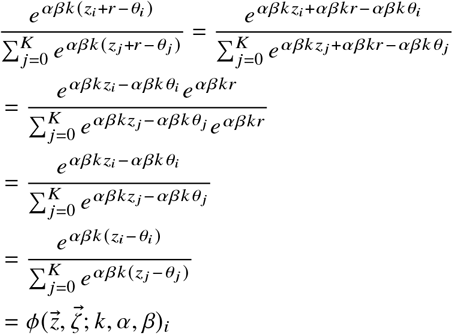

whereas for the Hill function:

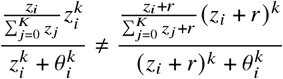

Thus the softmax function is less sensitive to sudden increases in all of the cues.

## 4. Conclusion

Multiple task integration should be a necessary component of any response threshold model which looks to explain the emergence of division of labor in social insect colonies. Tasks can sometimes be encountered sequentially if a task cue is locally bounded (such as wall building, Pinter-Wollman et al., 2012), but these insects can also be exposed to these tasks concurrently. For instance, both temperature and barometric pressure are felt throughout the nest (Abou-Shaara et al., 2017; Sujimoto et al., 2020) and alarm pheromone can interrupt the performance of a task (Sasaki et al., 2014; Guo et al., 2022). These coincident signals can have nonlinear effects on behavior. For instance, bees which perform a waggle dance on empty cells recruit more foragers than those that perform the dance on capped brood cells (Tautz, 1996). Additionally, as the nature of these cues are also not always known (Beshers & Fewell, 2001), modeling them as solely positive numbers may not always be appropriate. For example, the distance from the nest entrance on a return journey can be negative if an ant walks past the entrance (Wehner, 2020). Any comprehensive model of division of labor should account for the integration of simultaneous task cues and should also be able to handle negative cues. A Boltzmann sampler version of a response threshold is a strong candidate for such a model.

Consider Smith (2022) as an example for the importance of incorporating both negative cues and multiple tasks. Here, authors used the softmax function to model macronutrient regulation experiment in leafcutter ants. In this experiment, a colony could forage from one of two food sources with varying amounts of protein and carbohydrates. One food item had a higher carbohydrate to protein ratio than the colony’s intake target, while the other had a lower ratio, so the colony would need to collect both to properly regulate the growth of its fungal gardens. Authors expected that specialists would arise in this system, where the number of foraging trips per ant would be correlated with the proportion of trips to one food item or the other. No such correlation arose. Instead, there seemed to be no relationship between the number of trips a forger took and the proportion of trips to one food item or the other. In the Boltzmann model of this experiment, the distance between the colony’s current nutritional status and the intake target is the task-associated cue, and this distance can be either positive or negative. The probability of foraging from one food source depends just on this cue, while the probability of foraging from the other depends on the negative cue. Each ant also has two fixed, independently drawn response thresholds, one for each food item. Simulations recreated seemingly random experimental distributions of behavior, showing that the number of foraging trips was a result of the sum of the thresholds, whereas the propensity to choose one food item over the other was determined by the difference in thresholds. Thus, a threshold model can potentially describe a biological phenomenon, even in the absence of apparent specialists. This model, like others before it (Ulrich et al., 2021) shows the importance of considering multiple thresholds simultaneously, as well as the need to sometimes consider negative cues.

The distance from an ideal intermediate value does not necessarily have to be negative, but there are issues with alternative approaches. Consider Gove et al. (2009), where individuals integrate task cues with a two-step process. First, they take the absolute value of the difference between the cue and its optimal value. If this difference is greater than the response threshold (and it’s greater than the differences that correspond to other tasks), the individual will then check if the cue is higher or lower than its optimal value. If it’s higher, then the worker will perform the task that will lower the cue. If it’s lower, the worker will engage in the task which will raise the cue. The issue with this design is that it maps a single threshold onto two different tasks, implying that if she is a specialist in one task, she will also be a specialist in the other task, which is not necessarily the case (Weidenmüller, 2004). Additionally, in this model the cue is compared to the response threshold every timestep, so the worker will keep performing the task until the task is either completed or the need for another task overtakes it. This also implies a correlation between responsiveness and duration of task performance which also may not exist in real social insects (Weidenmüller, 2004).

This example highlights the need for the careful design of models which reflect biological reality, as ‘predictions’ of the model may not result from the response or satisfaction threshold concept directly, but rather from assumptions inadvertently built into the model. For instance, models which use the Hill function for the response threshold implicitly assume that the probability of starting a task is 0 when the task-associated cue is also 0, which seems like an overly strong assumption. Conversely, the probability of starting a task is never quite 1, introducing an asymmetry to the response curve which decreases the probability that the colony will respond to the task and subsequently raising the equilibrium values of the cue. The Hill function also introduces an inverse relationship between the gradient of the probability curve and the threshold, rather than having these features be controlled by two independent parameters. Finally, multi-task generalizations of the Hill function assume that when the cues for each task are high, the differences between them are relatively small, so the probability of doing each will be equivalent regardless of the relative values of the response or satisfaction thresholds.

The Boltzmann sampler makes none of these assumptions, but it is flexible enough to be remolded so that it can take on these assumptions should there be biological motivation to do so. For example, the Boltzmann sampler is scale-free, meaning that a simulation can be ‘zeroed’ at any value of the cue and the workers will behave in the same way. The same is not true of models which use the Hill function, where the probabilities of performing tasks becomes more uniform if the cue is much higher than the thresholds. A way to interpret this result for the Hill case is to say that when several cues are saturating the antenna, then the ant loses the ability to discriminate between the tasks and chooses to do one task randomly. This could be biologically reasonable, but this situation can also be modeled in the softmax case, where the steepness parameter could be turned into a function of the cues. Lowering the steepness parameter can make these probabilities more uniform for the softmax function, so one can imagine devising a function that lowers the steepness parameter when the ant is distracted by many cues. However, the Hill function is not so easily controlled. Even with high values of the steepness parameter and low values of the threshold, the probabilities of performing each task stay fairly uniform if the cues themselves are at similar levels. Thus, the softmax is the more flexible alternative to integrating multiple cues, as it can maintain a responsive system even with similar cues values while it can also be made to model scenarios where the social insect cannot distinguish between cues.

The Boltzmann sampler also allows for more control in the sense that the steepness of the probability curves is only determined by a single parameter; it does not implicitly assume a relationship between the two as the Hill function does. This conflation can make it difficult to interpret the effect of each of these parameters on a model in a sensitivity analysis, and it limits the number of potential configurations. The effects of the steepness and threshold parameters are inversely proportional, one cannot have a steeply-curved probability distribution that also has a high threshold. As these two parameters are independent of each other in the softmax function, though, then any combination of parameters is possible there. Again though, the softmax function can be forced to operate like the Hill function if one chooses by setting the two to be inversely proportional, but the reverse is not true.

The independence between parameters and scale-free properties are also important because the variables used to represent cues in social insect response threshold models are unitless (Bonabeau et al., 1996; Jeanson et al., 2007; Ulrich et al., 2021; Lynch et al., *in submission*). For many tasks, it is difficult to know what the task-associated cue is, let alone how to measure it (Beshers & Fewell, 2001), and so levels of the cue are chosen arbitrarily for the model. These levels could artificially influence the performance of models using a Hill function, as high levels can homogenize probabilities and decrease the steepness of the probability curve. However, these issues are not present for the softmax model, and since it can also take on negative values, the cue can be set to any level without influencing performance.

Taken together, these properties of the Boltzmann sampler makes it a good extension of the Hill function while also improving on it, as workers operate more like one would expect given the verbal description of the response and satisfaction threshold hypotheses. The Boltzmann sampler gives the maximal entropy distribution of probabilities, which can be interpreted as the distribution of least assumption. Additionally, these themes of control should also make the softmax function an attractive option for a task allocation algorithm for swarm robotics, which already rely heavily on social insect inspired Hill functions (Ducatelle et al., 2010; Kanakia et al., 2016; Wu et al., 2018; Jiang et al., 2020; Obute et al., 2022).

## Supplemental Information

### Partial derivative of softmax function

Let 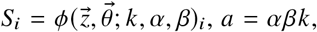, and 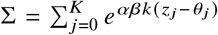. The partial derivative for the probability of performing task *i* with respect to the cue of an arbitrary *j* task is:

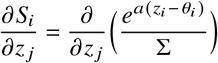

In the case where *i* = *j*:

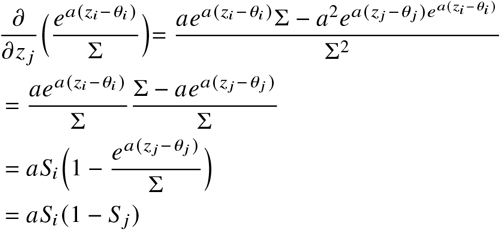

When *i* ≠ *j*:

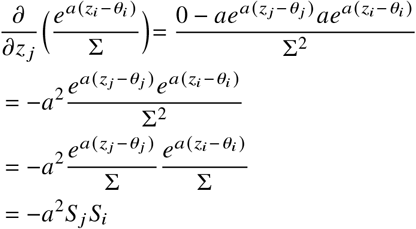

So in summary:

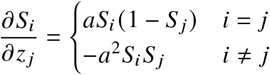

## Acknowledgments

We would also like to thank members of the Harrison and Fewell labs for comments made on the manuscript. We would also like to thank Michaela Starkey for her assistance in developing some mathematical formulations.

## References

Abou-Shaara, H. F., Owayss, A. A., Ibrahim, Y. Y., & Basuny, N. K. (2017). A review of impacts of temperature and relative humidity on various activities of honey bees. Insectes Sociaux, 64(4), 455–463.

Arcuri, A., & Lanchier, N. (2017). Stochastic spatial model for the division of labor in social insects. Mathematical Models and Methods in Applied Sciences, 27(01), 45–73.

Beshers, S. N., & Fewell, J. H. (2001). Models of division of labor in social insects. Annual review of Entomology, 46, 413.

Beshers, S. N., Robinson, G. E., & Mittenthal, J. E. (1999). Response thresholds and division of labor in insect colonies. In Information processing in social insects (pp. 115–139). Birkhäuser, Basel.

Bonabeau, E., Theraulaz, G., & Deneubourg, J. L. (1996). Quantitative study of the fixed threshold model for the regulation of division of labour in insect societies. Proceedings of the Royal Society of London. Series B: Biological Sciences, 263(1376), 1565–1569.

Bouyé, E., Durrleman, V., Nikeghbali, A., Riboulet, G., & Roncalli, T. (2000). Copulas for finance-a reading guide and some applications. Available at SSRN 1032533.

Calabi, P. (1988). Behavioral flexibility in Hymenoptera: a re-examination of the concept of caste. In Advances in Myrmecology (ed. J. C. Trager), pp. 237–258. E.J. Brill, New York.

Camazine, S., Deneubourg, J. L., Franks, N. R., Sneyd, J., Theraulaz, G. & Bonabeau, E. (2001). Self-Organization in Biological Systems. Princeton University Press, Princeton.

Dornhaus, A., Leitner, N., Lynch, N., Mallmann-Trenn, F., & Pajak, D. (2019). Remember the Past and Forget Thresholds. In 7th Workshop on Biological Distributed Algorithms (BDA).

Ducatelle, F., Di Caro, G. A., & Gambardella, L. M. (2010). Cooperative self-organization in a heterogeneous swarm robotic system. In Proceedings of the 12th annual conference on Genetic and Evolutionary Computation (pp. 87–94).

Duchon, P., Flajolet, P., Louchard, G., & Schaeffer, G. (2004). Boltzmann samplers for the random generation of combinatorial structures. Combinatorics, Probability and Computing, 13(4-5), 577–625.

Dussutour, A., Nicolis, S. C., Shephard, G., Beekman, M., & Sumpter, D. J. (2009). The role of multiple pheromones in food recruitment by ants. Journal of Experimental Biology, 212(15), 2337–2348.

Garrison, L. K., Kleineidam, C. J., & Weidenmüller, A. (2018). Behavioral flexibility promotes collective consistency in a social insect. Scientific Reports, 8(1), 15836.

Giurfa, M., & Sandoz, J. C. (2012). Invertebrate learning and memory: fifty years of ol-factory conditioning of the proboscis extension response in honeybees. Learning & memory, 19(2), 54–66.

Gove, R., Hayworth, M., Chhetri, M., & Rueppell, O. (2009). Division of labour and social insect colony performance in relation to task and mating number under two alternative response threshold models. Insectes sociaux, 56, 319–331.

Guo, X., Lin, M. R., Azizi, A., Saldyt, L. P., Kang, Y., Pavlic, T. P., & Fewell, J. H. (2022). Decoding alarm signal propagation of seed-harvester ants using automated movement tracking and supervised machine learning. Proceedings of the Royal Society B, 289(1967), 20212176.

Feng, T., Charbonneau, D., Qiu, Z., & Kang, Y. (2021). Dynamics of task allocation in social insect colonies: scaling effects of colony size versus work activities. Journal of Mathematical Biology, 82(5), 1–53.

Fewell JH, Page RE. (1999) The emergence of division of labour in forced associations of normally solitary ant queens. Evol Ecol Res. 1:537–48.

Holbrook, C. T., Barden, P. M., & Fewell, J. H. (2011). Division of labor increases with colony size in the harvester ant Pogonomyrmex californicus. Behavioral Ecology, 22(5), 960–966.

Holbrook CT, Kukuk PF, Fewell JH. (2013) Increased group size promotes task specialization in a normally solitary halictine bee. Behaviour. 150:1449–66.

Jeanson, R., Fewell, J. H., Gorelick, R., & Bertram, S. M. (2007). Emergence of increased division of labor as a function of group size. Behavioral Ecology and Sociobiology, 62(2), 289–298.

Jeanson, R., & Lachaud, J. P. (2015). Influence of task switching costs on colony home-ostasis. The Science of Nature, 102(5), 1–4.

Jiang, C., Chen, T., Li, R., Li, L., Li, G., Xu, C., & Li, S. (2020). Construction of extended ant colony labor division model for traffic signal timing and its application in mixed traffic flow model of single intersection. Concurrency and Computation: Practice and Experience, 32(7), e5592.

Kanakia, A., Klingner, J., & Correll, N. (2016). A response threshold sigmoid function model for swarm robot collaboration. In Distributed Autonomous Robotic Systems (pp. 193–206). Springer, Tokyo.

Leitner, N., Lynch, C., & Dornhaus, A. (2019). Ants in isolation: obstacles to testing worker responses to task stimuli outside of the colony context. Insectes Sociaux, 66(3), 343–354.

Lin, M. R. (2021). Energy Use Scaling and Alarm Spread in Social Ants: An Investigation Using Multi-agent Simulation and Object Tracking (Doctoral dissertation, Arizona State University).

Lynch, C., Dornhaus, A, Wilson, R. New version of an old mechanism for task allocation in social insects. *In submission*.

Maccaro, J. J., Whyte, B. A., & Tsutsui, N. D. (2020). The ant who cried wolf? Short-term repeated exposure to alarm pheromone reduces behavioral response in Argentine ants. Insects, 11(12), 871.

Moreno, E., & Arenas, A. (2023). Changes in resource perception throughout the foraging visit contribute to task specialization in the honey bee Apis mellifera. Scientific Reports, 13(1), 8164.

Obute, S. O., Kilby, P., Dogar, M. R., & Boyle, J. H. (2022). Swarm Foraging Under Communication and Vision Uncertainties. IEEE Transactions on Automation Science and Engineering.

O’Donnell, S., & Foster, R. L. (2001). Thresholds of response in nest thermoregulation by worker bumble bees, Bombus bifarius nearcticus (Hymenoptera: Apidae). Ethology, 107(5), 387–399.

Pankiw, T., Page Jr, R. E., & Kim Fondrk, M. (1998). Brood pheromone stimulates pollen foraging in honey bees (Apis mellifera). Behavioral ecology and sociobiology, 44(3), 193–198.

Pankiw T, Page RE. (2000) Response thresholds to sucrose predict foraging division of labor in honeybees. Behav Ecol Sociobiol. 47:265–7.

Pinter-Wollman, N., Hubler, J., Holley, J. A., Franks, N. R., & Dornhaus, A. (2012). How is activity distributed among and within tasks in Temnothorax ants? Behavioral Ecology and Sociobiology, 66(10), 1407–1420.

Rajagopal, S., Brockmann, A., & George, E. A. (2022). Environment-dependent benefits of interindividual variation in honey bee recruitment. Animal Behaviour, 192, 9–26.

Ravary, F., Lecoutey, E., Kaminski, G., Châline, N., & Jaisson, P. (2007). Individual experience alone can generate lasting division of labor in ants. Current Biology, 17(15), 1308–1312.

Sadeh, S., & Clopath, C. (2022). Contribution of behavioural variability to representational drift. bioRxiv.

Sasaki, T., Hölldobler, B., Millar, J. G., & Pratt, S. C. (2014). A context-dependent alarm signal in the ant Temnothorax rugatulus. Journal of Experimental Biology, 217(18), 3229–3236.

Smith, N. (2022). Macronutrient Regulation by the Desert Leafcutter Ant Acromyrmex versicolor (Doctoral dissertation, Arizona State University).

Sujimoto, F. R., Costa, C. M., Zitelli, C. H., & Bento, J. M. S. (2020). Foraging activity of leaf-cutter ants is affected by barometric pressure. Ethology, 126(3), 290–296.

Sutton, R. S. and Barto A. G. (1998) Reinforcement Learning: An Introduction. The MIT Press, Cambridge, MA. Softmax Action Selection

Tautz, J. (1996). Honeybee waggle dance: recruitment success depends on the dance floor. The Journal of Experimental Biology, 199(6), 1375–1381.

Theraulaz, G., Bonabeau, E., & Denuebourg, J. N. (1998). Response threshold reinforcements and division of labour in insect societies. Proceedings of the Royal Society of London. Series B: Biological Sciences, 265(1393), 327–332.

Tschinkel, W. R. (2005). The nest architecture of the ant, Camponotus socius. Journal of Insect Science, 5(1), 9.

Ulrich Y, Saragosti J, Tokita CK, Tarnita CE, Kronauer DJC. (2018) Fitness benefits and emergent division of labor at the onset of group-living. Nature. 560:635–8. pmid:30135576

Ulrich, Y., Kawakatsu, M., Tokita, C. K., Saragosti, J., Chandra, V., Tarnita, C. E., & Kronauer, D. J. (2021). Response thresholds alone cannot explain empirical patterns of division of labor in social insects. PLoS Biology, 19(6), e3001269.

Wehner, R. (2020). Desert navigator. In Desert Navigator. Harvard University Press.

Weidenmüller, A. (2004). The control of nest climate in bumblebee (Bombus terrestris) colonies: interindividual variability and self reinforcement in fanning response. Behavioral Ecology, 15(1), 120–128.

Wu, H., Li, H., Xiao, R., & Liu, J. (2018). Modeling and simulation of dynamic ant colony’s labor division for task allocation of UAV swarm. Physica A: Statistical Mechanics and its Applications, 491, 127–141.

Yang, Y., Zhou, C., & Tian, Y. (2009). Swarm robots task allocation based on response threshold model. In 2009 4th International Conference on Autonomous Robots and Agents (pp. 171–176). IEEE.

